# Increased receptor binding capability of the SARS-CoV-2 saltational variant PJ.2.1

**DOI:** 10.64898/2026.09.03.749172

**Authors:** PeiZhuo He, BingKun Li, Caiwan Guo, Lingling Yu, Yuanling Yu, Fanchong Jian, Fei Shao, Yunlong Cao

**Affiliations:** Changping Laboratory, Beijing, P. R. China; Chinese Academy of Medical Sciences & Peking Union Medical College.; Biomedical Pioneering Innovation Center (BIOPIC), School of Life Sciences, Peking University, Beijing, P. R. China; Academy for Advanced Interdisciplinary Studies, Peking University, Beijing, P. R. China; Peking–Tsinghua Center for Life Sciences, Peking University, Beijing, P.R. China

## Abstract

The recently identified SARS-CoV-2 saltational variant PJ.2.1, a highly mutated descendant of MC.10.1, has achieved rapid intercontinental spread since its detection in May 2026. Here, we show that PJ.2.1 exhibits exceptionally high human ACE2-binding capability, significantly outperforming contemporary variants such as NB.1.8.1, XFG, and BA.3.2.2. However, despite acquiring over 25 spike mutations, PJ.2.1 remains antigenically similar to the circulating JN.1 family and does not display strong humoral immune evasion. Neutralization profiling indicates that PJ.2.1 possesses heightened neutralization sensitivity to human plasma and RBD-targeting antibodies, especially against cryptic-site-targeting class 4 and class 5 antibodies. This distinct sensitivity profile strongly suggests an altered spike structural dynamic that favors a receptor-accessible "up" conformation. While PJ.2.1 currently lacks the extreme immune evasion capabilities of other contemporary strains, its robust baseline receptor binding strength mirrors the early evolutionary trajectory of BA.2.86 to JN.1. This high receptor binding affinity provides a structural buffer that could facilitate the rapid acquisition of potent immune-evasive mutations, necessitating continued genomic, epidemiological, and virological surveillance of PJ.2.1.

## Results

The emergence of saltational SARS-CoV-2 variants, such as BA.1 and BA.2.86, has historically enhanced immune evasion, repeatedly reshaped the pandemic landscape, and prompted critical updates to vaccine formulations ^1–2^. As a highly mutated saltational descendant of MC.10.1, the recently discovered PJ.2.1 has acquired more than 25 spike mutations and achieved intercontinental spread. This raises global concerns regarding its potential to alter the SARS-CoV-2 epidemiological trajectory and impact future vaccine strain selection (Figure 1A). These factors warrant immediate investigation into the virological impact of the spike mutations on PJ.2.1.

**Figure 1.**
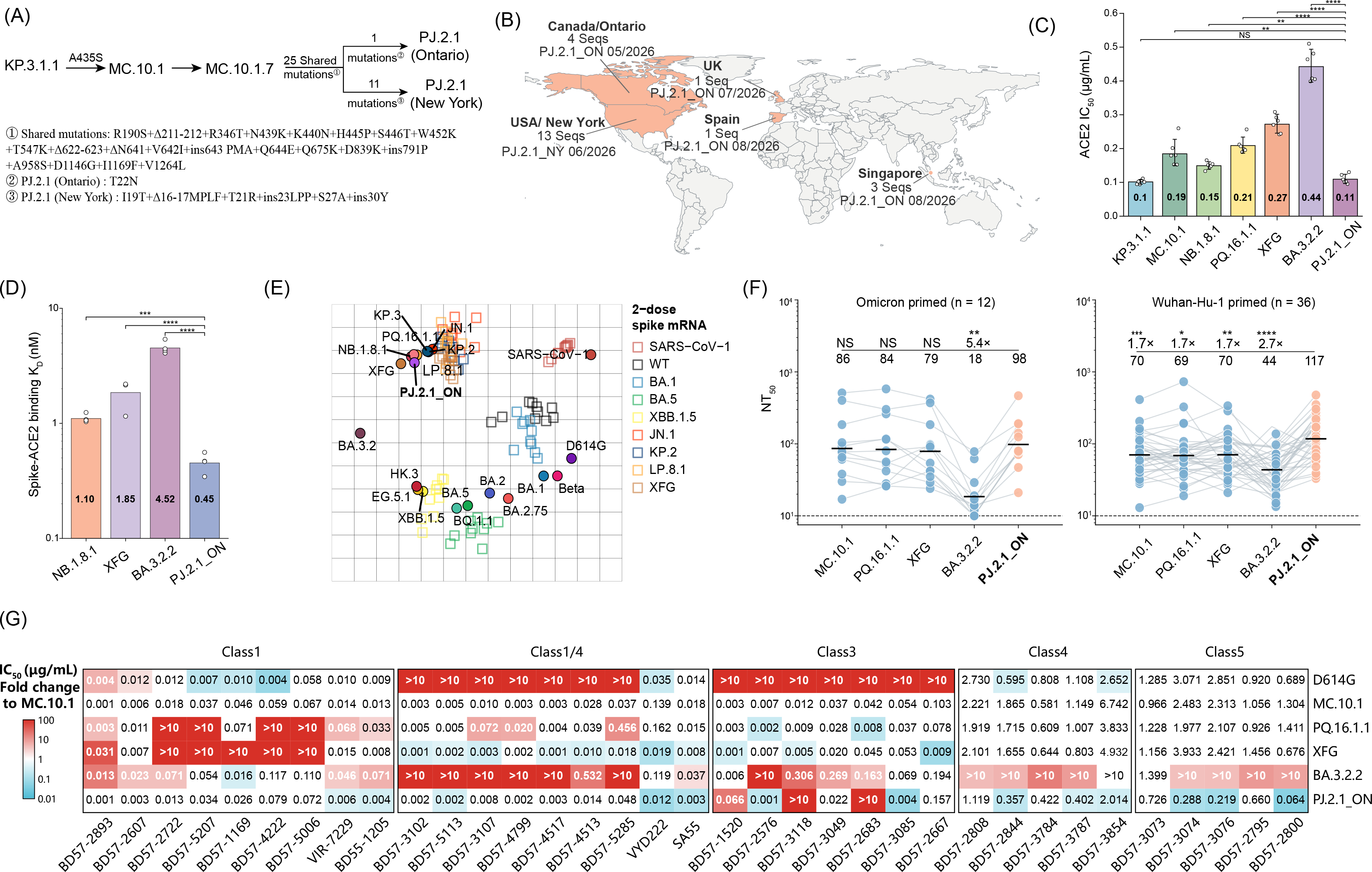
Virological, epidemiological, and antigenic characterization of the the SARS-CoV-2 saltation variant PJ.2.1. (A) Schematic evolutionary pathway and spike glycoprotein mutational profile of SARS-CoV-2 saltation variant PJ.2.1. PJ.2.1 (Ontario) and PJ.2.1 (New York) represent distinct clinical isolates. (B) Global geographic distribution of SARS-CoV-2 variant PJ.2.1. Coral shading highlights countries with detected cases. Labels indicate geographic location, sequence counts, corresponding isolate (PJ.2.1_ON or PJ.2.1_NY), and earliest collection dates (month/year), using data retrieved from GISAID and NCBI GenBank. (C) Soluble human ACE2 neutralization against KP.3.1.1, MC.10.1, NB.1.8.1, PQ.16.1.1, XFG, BA.3.2.2, and PJ.2.1_ON pseudoviruses. Columns and numeric values indicate geometric mean half-maximal inhibitory concentration (IC_50_, μg/mL) values, with scatter points representing independent technical replicates. Statistical significance was evaluated using a two-tailed t test on log-transformed IC_50_ values. (D) Surface plasmon resonance (SPR) measurement of human ACE2 (hACE2) binding affinities for NB.1.8.1, XFG, BA.3.2.2, and PJ.2.1_ON spike trimers. Bar heights and embedded values indicate geometric mean dissociation constants (K_D_, nM), with scatter points denoting individual technical replicates. Statistical significance was evaluated using two-tailed t test on log-transformed K_D_ values. (E) Antigenic cartography based on serum NT50 titers from mice immunized with two doses of spike mRNA vaccines against indicated variants. (F) Plasma neutralization titers against MC.10.1, PQ.16.1.1, XFG, BA.3.2.2, and PJ.2.1_ON pseudoviruses in Omicron-primed (n=12) and Wuhan-Hu-1-primed (n=36) cohorts. Solid horizontal lines and numerical values designate geometric mean titers (NT_50_), with individual connected dots indicating paired plasma samples. The horizontal dashed line marks the limit of detection (NT_50_=10). Statistical comparisons, fold changes relative to PJ.2.1_ON, and significance levels assessed by the Wilcoxon rank-sum test are annotated above each variant. (G) Neutralization profiles of RBD-directed monoclonal antibodies across distinct epitope classes against indicated SARS-CoV-2 pseudoviruses. Background color gradient depicts fold changes relative to MC.10.1, where red indicates increased resistance and blue denotes heightened sensitivity according to the color scale. IC_50_=50% inhibitory concentration. NS=not significant. NT_50_=50% neutralisation titre. *p<0.05. **p<0.01. ***p<0.001. ****p<0.0001.

The first PJ.2.1 sequence was detected in Ontario, Canada, in May 2026. Subsequent wastewater surveillance identified the variant in four New York City sewersheds and one in Utah. Clinical cases quickly emerged in June 2026, followed by international reports from the UK, Spain, and Singapore, underscoring its rapid geographic spread (Figure 1B). Analysis of sequences deposited in the GISAID EpiCoV database revealed two distinct PJ.2.1 lineages that diverge primarily based on N-terminal domain (NTD) mutations. We designated these as PJ.2.1_ON (Ontario) and PJ.2.1_NY (New York) (Figure 1A). Because PJ.2.1_ON was identified earlier and shares key sequence characteristics with the clinical cases reported internationally, we focused our current investigation on this specific lineage.

PJ.2.1 harbors multiple substitutions within the receptor-binding domain (RBD) on sites previously frequently observed mutating, including R346T, N439K, K440N, 445P, 452K, and S446T. These substitutions, thoroughly documented in prior deep mutational scanning and experimental studies, are known to modulate ACE2 binding affinity and mediate antibody escape ^3–5^. Additionally, PJ.2.1 has acquired more than 10 mutations across the SD1, 630-loop region, S1/S2 cleavage sites, and S2 subunit domain. This extensive mutational profile indicates a high probability of significant alterations in the structural configuration of the spike protein. To assess whether PJ.2.1 possesses virological and antigenic characteristics associated with a potential transmission advantage, we compared it with its ancestral strain, MC.10.1 (a sublineage of KP.3.1.1), as well as the currently circulating variants NB.1.8.1, PQ.16.1.1, BA.3.2.2, and XFG ^6–8^.

We first generated spike-pseudotyped vesicular stomatitis viruses bearing the spike proteins of PJ.2.1_ON, MC.10.1, KP.3.1.1, NB.1.8.1, PQ.16.1.1, and XFG, assessing their receptor engagement capability via susceptibility to inhibition by soluble human angiotensin-converting enzyme 2 (hACE2). PJ.2.1_ON exhibited an hACE2 inhibition IC50 comparable to that of KP.3.1.1 and slightly lower than that of MC.10.1. Crucially, its IC50 was substantially lower than those of the contemporary circulating variants NB.1.8.1, PQ.16.1.1, and XFG (Figure 1C). Because a lower hACE2 inhibition IC_50_ indicates more efficient interaction with soluble hACE2, these findings suggest that PJ.2.1_ON has a higher hACE2-binding affinity and more efficient receptor engagement than contemporary lineages.

Next, we generated stabilized spike trimers (S6P) corresponding to NB.1.8.1, XFG, BA.3.2.2, and PJ.2.1_ON to quantify their binding affinities for soluble hACE2 utilizing surface plasmon resonance (SPR). Among the tested variants, PJ.2.1_ON exhibited the highest hACE2-binding affinity, followed by NB.1.8.1, XFG, and BA.3.2.2. These results are consistent with the hACE2 inhibition assay, directly indicating the superior hACE2 engagement by PJ.2.1_ON compared to the other tested variants (Figure 1D and S1). Collectively, these findings suggest the unique set of mutations in PJ.2.1_ON enhances the spike-ACE2 interaction, potentially facilitating more efficient cellular entry. This enhanced receptor engagement is likely driven not solely by RBD mutations, but also by NTD, 630-loop, and S1/S2 cleavage site substitutions, which exert allosteric effects that may alter the RBD “up” and “down” conformational dynamics.

We also evaluated the antigenic properties and humoral immune evasion profiles of these variants using pseudovirus neutralization assays. Antigenicity was mapped using sera from naive mice immunized twice with spike-encoding mRNA vaccines, while humoral immune evasion was evaluated using recent plasma sourced from humans with varying exposure histories. Antigenic cartography based on mice sera revealed that PJ.2.1 surprisingly clustered closely with members of the currently circulating JN.1 family, indicating relatively short antigenic distances (Figure 1E). In plasma from Omicron-primed individuals (defined as unvaccinated cohorts with a primary Omicron infection history), the neutralization sensitivity of PJ.2.1_ON was comparable to MC.10.1, KP.3.1.1, NB.1.8.1, and XFG, with no statistically significant differences in neutralization titers (Figure 1F). In contrast, in plasma from individuals primed with inactivated Wuhan-Hu-1 vaccines, neutralization titers against PJ.2.1_ON were even significantly higher than those against the other variants tested, including MC.10.1 (Figure 1F). These results confirm that PJ.2.1_ON remains antigenically similar to circulating JN.1-family variants, has not developed substantial humoral immune evasion capabilities, and may even display increased susceptibility to antibody neutralization.

We further extended our immunological profiling utilizing a comprehensive panel of RBD-targeting neutralizing monoclonal antibodies (mAbs) representing distinct epitope classes ^4–5^. Compared to MC.10.1, PJ.2.1_ON did not exhibit increased evasion against class 1 and class 1/4 antibodies (Figure 1G). This contrasts with XFG and PQ.16.1.1, which demonstrated substantially greater immune evasion against class 1 mAbs, and BA.3.2.2, which exhibited marked evasion against class 1/4 mAbs relative to PJ.2.1_ON. However, likely due to multiple mutations within the 439–450 loop, PJ.2.1_ON demonstrated a BA.3.2.2-like enhanced resistance to class 3 antibodies. Notably, these class 3 antibodies are largely Omicron-specific and are predominantly elicited in Omicron- primed individuals or through Omicron-induced de novo antibody responses ^9^.

Most importantly, PJ.2.1_ON displayed increased neutralization sensitivity compared to the other tested variants when exposed to class 4 and class 5 antibodies (Figure 1G). These mAbs target cryptic sites on the RBD that remain sterically inaccessible within a fully closed (3-down) spike trimer conformation. This contrast is especially pronounced when comparing PJ.2.1_ON with BA.3.2.2; the latter predominantly adopts an enriched 3-down RBD structural configuration and consequently showed minimal neutralization by cryptic-site-targeting mAbs ^10^. The heightened neutralization sensitivity of PJ.2.1_ON—observed not only against cryptic-site targeting mAbs but also against class 1 mAbs, class 1/4 mAbs, and human plasma—strongly suggests an altered RBD conformational dynamic that favors the receptor-accessible “up” state, thereby exposing the internal epitopes targeted by class 4 and class 5 antibodies and improving accessibility for “up” RBD- targeting neutralization mAbs.

Taken together, we found that while PJ.2.1 lacks the extreme immune evasion capabilities of past wave drivers, it possesses a significantly higher human ACE2 binding affinity than currently dominant variants. This distinct phenotypic profile suggests PJ.2.1 may follow an evolutionary trajectory analogous to BA.2.86/JN.1. BA.2.86 similarly leveraged a high-affinity structural foundation to subsequently acquire the L455S mutation, ultimately evolving into the highly evasive JN.1 variant that drove widespread global infections in 2024. Their robust ACE2 binding provides a structural buffer, enabling the rapid acquisition of strong immune-evasive mutations at the expense of binding affinity. Although current findings do not definitively establish an immediate transmission advantage, the global reporting of PJ.2.1 highlights the critical risk of further adaptation. The potential acquisition of known immune evasion-enhancing substitutions—such as A475V, N487D, and D420N, which are frequently observed in recent variants—could immediately enable PJ.2.1 descendants to reshape the epidemiological landscape. Consequently, continued genomic, epidemiological, and virological surveillance of PJ.2.1 is essential.

## Appendix

### Acknowledgments

We extend our gratitude to the scientific community for their continued efforts in monitoring SARS-CoV-2 variants, as well as to all volunteers who contributed blood samples for this study. This project is financially supported by Changping Laboratory (2026D-04-01 to Y.C.).

### Author Contributions

Y.C. designed and supervised the study. P.H., B.L. and Y.C. wrote the manuscript with input from all authors. P.H., B.L. and C.G. performed the sequence analyses and prepared the illustrations. Y.Y. constructed the pseudoviruses. L.Y. and F.S. processed the plasma samples and performed the pseudovirus neutralisation assays. P.H. and Y.C. analysed the neutralisation data.

## Methods Details

### Genomic sequence analysis and lineage definition

SARS-CoV-2 sequence metadata for the saltation variant PJ.2.1 and related reference lineages evaluated in this study were retrieved from the GISAID EpiCoV^1^ and NCBI GenBank databases on September 1, 2026. For the emerging saltation variant PJ.2.1, all available records collected from May to August 2026. Lineage assignments and aliases were verified according to the Pango nomenclature criteria outlined in lineage_notes.txt and alias_key.json from the cov-lineages/pango- designation repository. PJ.2.1 isolates identified from Ontario (Canada) and New York (USA) were referred to as PJ.2.1_ON and PJ.2.1_NY. These curated data were used to determine the geographic distribution, total sequence counts, and earliest collection dates across affected countries.

### Surface Plasmon Resonance

Binding kinetics of SARS-CoV-2 spike trimers to human ACE2 were determined on a Biacore 8K instrument (Cytiva, cat. no. 12914229). Human ACE2 was captured onto Protein A sensor chips (Cytiva, cat. no. 29127556). Purified spike trimer ectodomains of the indicated variants were prepared in a twofold serial dilution series across six concentrations (ranging from 3.125 to 100 nM) and flowed across the chip surface in multi-cycle kinetics mode at ambient temperature. Sensorgrams were acquired with Biacore 8K Control Software (v.4.0.8.19879; Cytiva). Kinetic association and dissociation rates as well as equilibrium dissociation constants (K_D_) were obtained through global fitting to a 1:1 binding model using Biacore 8K Evaluation Software (v.4.0.8.20368; Cytiva).

### Cohort recruitment and plasma processing

Peripheral blood samples were obtained from individuals enrolled in the study following the acquisition of written informed consent. Consent covered specimen procurement, biorepository storage, scientific investigation, and subsequent data dissemination. Study protocols received formal approval from the Tianjin Municipal Health Commission, the Institutional Review Board of Tianjin First Central Hospital (approval no. KEYAN20241022-2), and the Ethics Committee of Beijing Ditan Hospital, Capital Medical University (approval no. DTEC-KY2024-112-01). All procedures complied strictly with the ethical standards of the Declaration of Helsinki.For specimen isolation, whole blood was mixed 1:1 (v/v) with phosphate-buffered saline supplemented with 2% fetal bovine serum. Density-gradient centrifugation using Ficoll-Paque media (Cytiva, cat. no. 17-1440-03) was conducted to fractionate peripheral blood mononuclear cells and plasma. The cell-free plasma supernatant was harvested, divided into single-use aliquots, and cryopreserved at – 20° C or lower. Prior to downstream neutralization assays, all plasma specimens were heat-inactivated at 56°C for 30 min.

### Pseudovirus neutralization assay

SARS-CoV-2 spike pseudoviruses were produced using a vesicular stomatitis virus (VSV)-based packaging system. Briefly, 293T cells (ATCC, CRL-3216) were transfected with spike-expression plasmids together with G*ΔG-VSV (Kerafast). Culture supernatants were harvested, filtered, aliquoted, and stored at −80°C.For neutralization assays, pseudoviruses were incubated with serially diluted monoclonal anti-S-RBD antibodies, human ACE2-Fc, or heat-inactivated plasma in 96-well plates at 37°C with 5% CO_2_ for 1 h. Plasma samples were initially diluted 1:10, followed by five consecutive threefold dilutions. Monoclonal antibodies and human ACE2-Fc (initial concentration 0.1 mg/mL) were pre-diluted 1:100 and 1:50, respectively, followed by five consecutive fivefold dilutions. Huh-7 cells (JCRB, 0403) were then added to each well. After 24 h of incubation, supernatants were removed and D-luciferin reagent (Vazyme, DD1209-03) was added. Luminescence was measured using a microplate spectrophotometer (PerkinElmer, HH3400) after 2 min in the dark. Half-maximal inhibitory concentration (IC_50_) and NT_50_ values were calculated using a four-parameter logistic regression model.

## Supplementary Tables

**Table S1.**
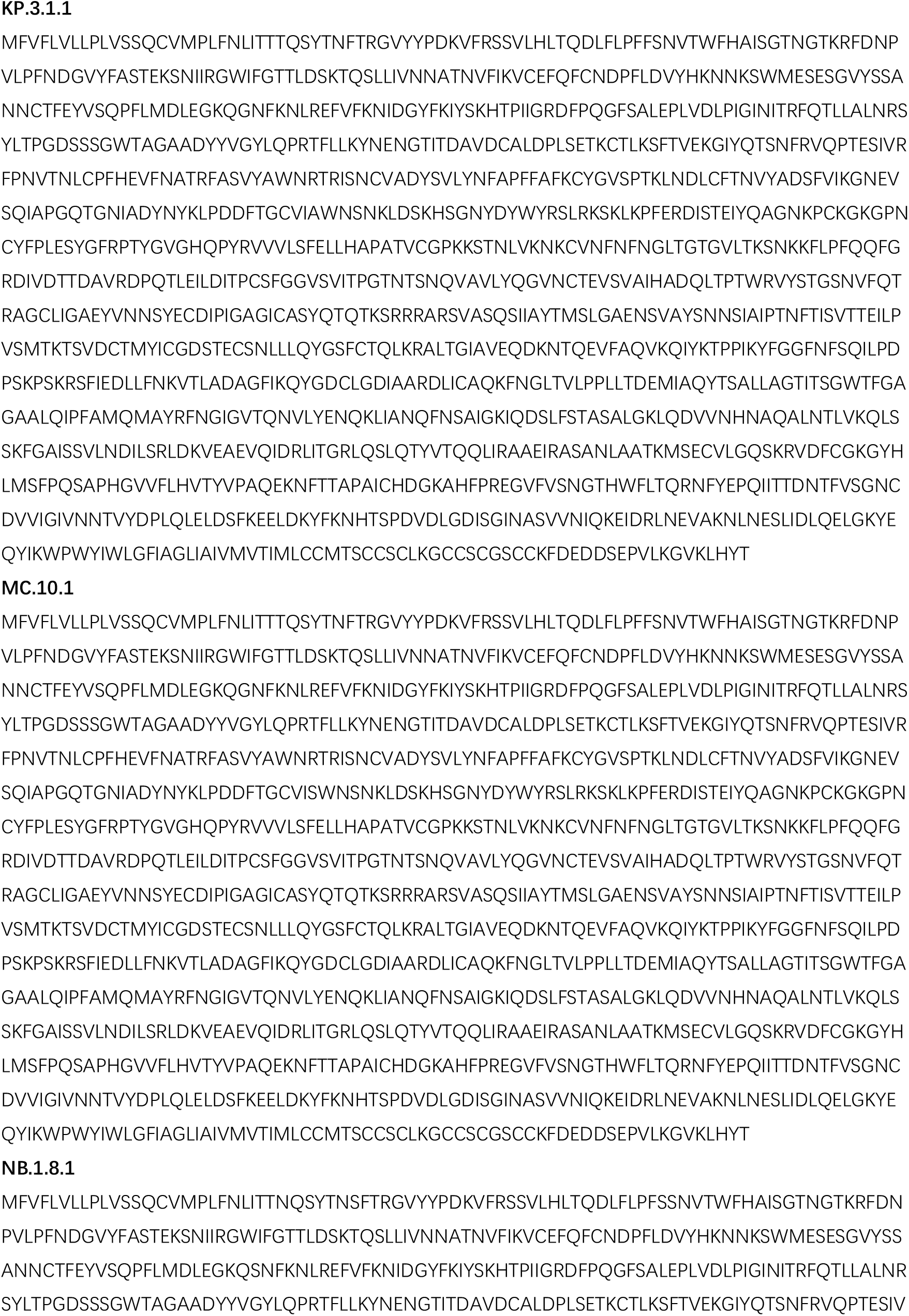
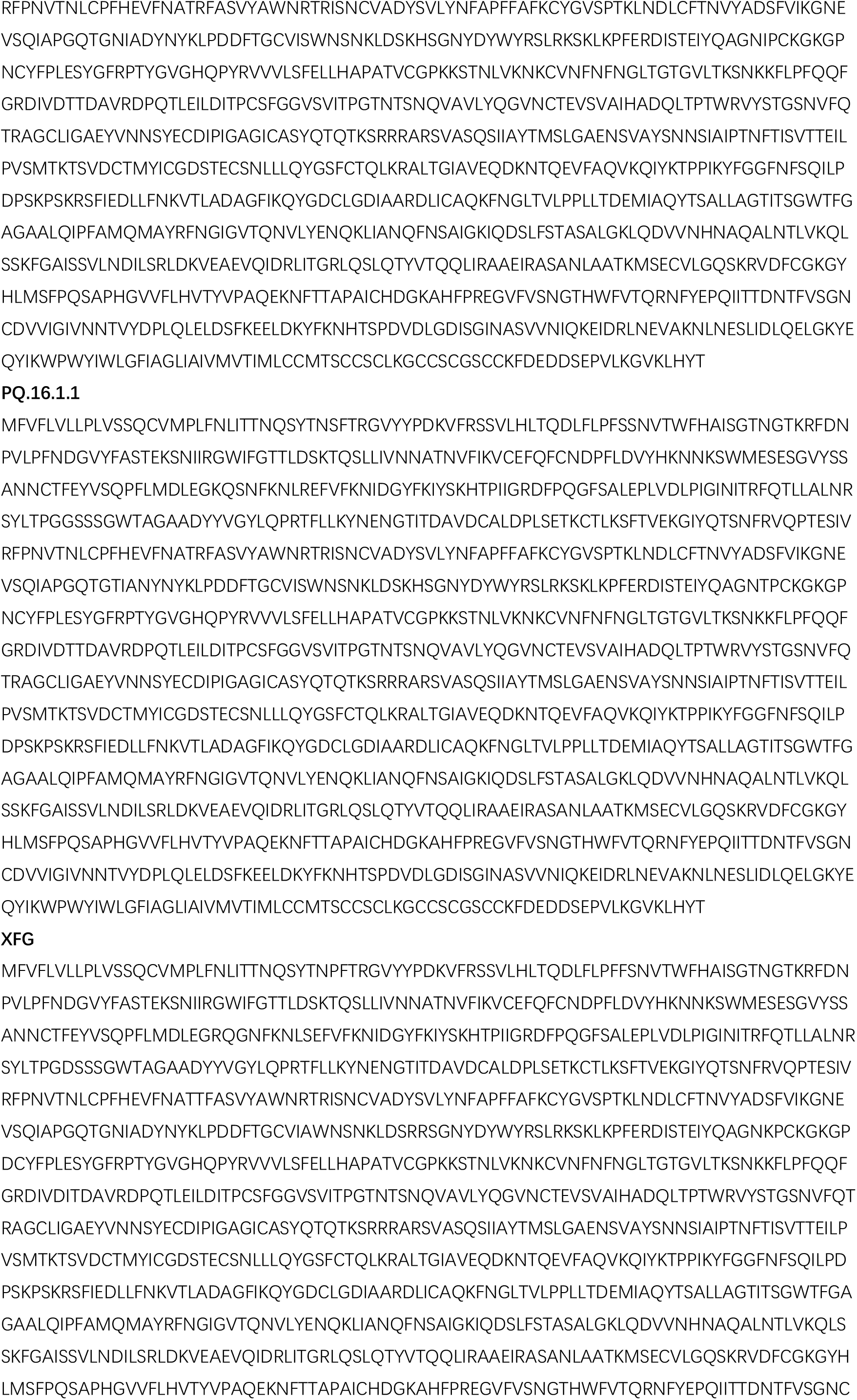
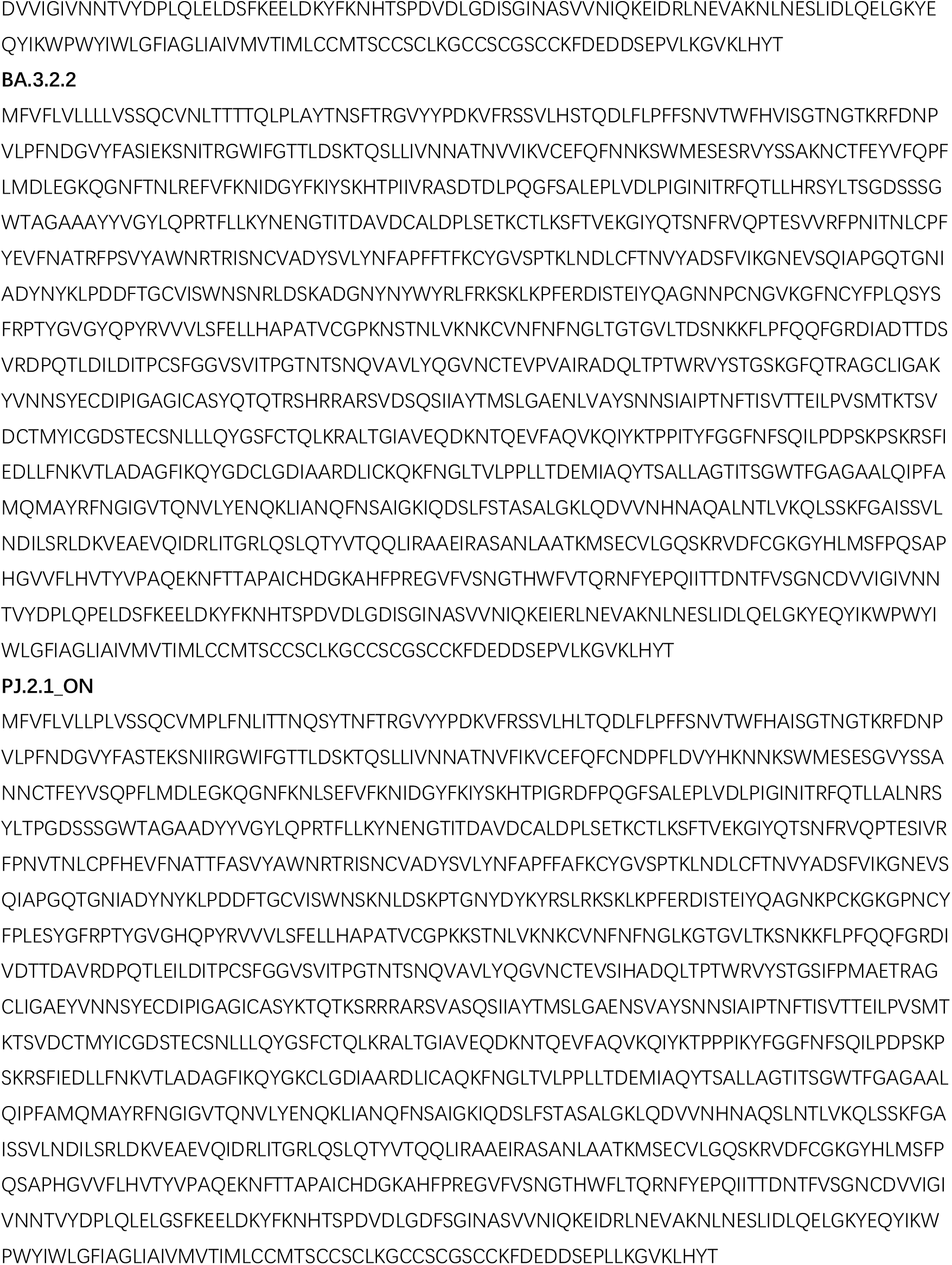
Spike protein sequences of SARS-CoV-2 variants.

**Table S2.** Kinetic parameters of SARS-CoV-2 variant spike binding to hACE2 measured by SPR.

| **Sample** | **$K_a(M^{-1} s^{-1})$** | **$K_d (s^{-1})$** | **$K_D(M)$** |
| --- | --- | --- | --- |
| NB.1.8.1-Spike | 1.11E+05 | 1.20E-04 | 1.08E-09 |
| NB.1.8.1-Spike | 1.07E+05 | 1.33E-04 | 1.24E-09 |
| NB.1.8.1-Spike | 1.12E+05 | 1.16E-04 | 1.03E-09 |
| NB.1.8.1-Spike | 1.07E+05 | 1.13E-04 | 1.06E-09 |
| XFG-Spike | 8.50E+04 | 1.84E-04 | 2.16E-09 |
| XFG-Spike | 8.64E+04 | 1.86E-04 | 2.15E-09 |
| XFG-Spike | 1.33E+05 | 1.53E-04 | 1.15E-09 |
| XFG-Spike | 7.92E+04 | 1.74E-04 | 2.19E-09 |
| PJ.2.1_ON-Spike | 2.88E+05 | 9.91E-05 | 3.44E-10 |
| PJ.2.1_ON-Spike | 1.81E+05 | 1.01E-04 | 5.59E-10 |
| PJ.2.1_ON-Spike | 1.96E+05 | 9.19E-05 | 4.68E-10 |
| PJ.2.1_ON-Spike | 1.96E+05 | 9.11E-05 | 4.66E-10 |
| BA.3.2.2-Spike | 4.54E+04 | 1.85E-04 | 4.08E-09 |
| BA.3.2.2-Spike | 4.22E+04 | 1.81E-04 | 4.29E-09 |
| BA.3.2.2-Spike | 4.02E+04 | 2.16E-04 | 5.38E-09 |
| BA.3.2.2-Spike | 4.35E+04 | 1.93E-04 | 4.44E-09 |

**Table S3.** Information of recruited SARS-CoV-2 convalescent participants.

| Sample Type | Gender | Age | Wuhan-1 primed | Sampling time point | NT<sub>50</sub> |  |  |  |  |
| --- | --- | --- | --- | --- | --- | --- | --- | --- | --- |
|  |  |  |  |  | MC.10.1 | PQ.16.1.1 | XFG | BA.3.2.2 | PJ.2.1 ON |
| Omicron primed individuals | female | 39 | no | 2025/4/9 | 510 | 580 | 419 | 62 | 466 |
| Omicron primed individuals | female | 47 | no | 2025/4/9 | 93 | 118 | 42 | 12 | 47 |
| Omicron primed individuals | male | 75 | no | 2025/5/14 | 30 | 28 | 98 | 15 | 76 |
| Omicron primed individuals | female | 38 | no | 2025/5/28 | 78 | 53 | 95 | 29 | 141 |
| Omicron primed individuals | female | 21 | no | 2025/6/26 | 104 | 116 | 54 | 19 | 71 |
| Omicron primed individuals | female | 42 | no | 2025/7/2 | 185 | 261 | 185 | 78 | 193 |
| Omicron primed individuals | female | 79 | no | 2025/7/16 | 77 | 80 | 91 | 10 | 77 |
| Omicron primed individuals | male | 51 | no | 2025/7/16 | 17 | 26 | 24 | 14 | 21 |
| Omicron primed individuals | female | 52 | no | 2025/7/24 | 38 | 26 | 42 | 10 | 81 |
| Omicron primed individuals | female | 42 | no | 2025/8/13 | 61 | 42 | 29 | 14 | 125 |
| Omicron primed individuals | female | 68 | no | 2025/8/13 | 402 | 301 | 372 | 17 | 190 |
| Omicron primed individuals | female | 43 | no | 2025/8/13 | 65 | 56 | 35 | 10 | 79 |
| Wuhan-Hu-1 primed individual | female | 28 | yes | 2025/4/9 | 80 | 86 | 78 | 60 | 101 |
| Wuhan-Hu-1 primed individual | female | 57 | yes | 2025/4/16 | 130 | 76 | 160 | 48 | 213 |
| Wuhan-Hu-1 primed individual | female | 55 | yes | 2025/4/16 | 65 | 55 | 134 | 57 | 108 |
| Wuhan-Hu-1 primed individual | female | 40 | yes | 2025/4/16 | 59 | 81 | 94 | 39 | 145 |
| Wuhan-Hu-1 primed individual | female | 45 | yes | 2025/4/16 | 146 | 108 | 105 | 89 | 477 |
| Wuhan-Hu-1 primed individual | female | 31 | yes | 2025/4/23 | 28 | 35 | 63 | 26 | 89 |
| Wuhan-Hu-1 primed individual | female | 25 | yes | 2025/4/17 | 47 | 63 | 58 | 45 | 68 |
| Wuhan-Hu-1 primed individual | male | 36 | yes | 2025/4/24 | 37 | 24 | 47 | 55 | 132 |
| Wuhan-Hu-1 primed individual | male | 28 | yes | 2025/4/24 | 48 | 91 | 27 | 20 | 33 |
| Wuhan-Hu-1 primed individual | female | 36 | yes | 2025/4/24 | 65 | 85 | 121 | 71 | 117 |
| Wuhan-Hu-1 primed individual | female | 56 | yes | 2025/4/24 | 38 | 56 | 102 | 21 | 104 |
| Wuhan-Hu-1 primed individual | male | 61 | yes | 2025/5/14 | 30 | 45 | 37 | 38 | 57 |
| Wuhan-Hu-1 primed individual | female | 39 | yes | 2025/5/14 | 412 | 730 | 287 | 85 | 351 |
| Wuhan-Hu-1 primed individual | female | 35 | yes | 2025/5/23 | 119 | 190 | 106 | 93 | 112 |
| Wuhan-Hu-1 primed individual | female | 31 | yes | 2025/5/15 | 32 | 67 | 36 | 28 | 36 |
| Wuhan-Hu-1 primed individual | female | 36 | yes | 2025/5/15 | 55 | 94 | 122 | 29 | 187 |
| Wuhan-Hu-1 primed individual | male | 27 | yes | 2025/5/15 | 49 | 59 | 52 | 79 | 156 |
| Wuhan-Hu-1 primed individual | female | 48 | yes | 2025/5/15 | 151 | 193 | 338 | 36 | 247 |
| Wuhan-Hu-1 primed individual | female | 75 | yes | 2025/5/28 | 206 | 326 | 43 | 174 | 84 |
| Wuhan-Hu-1 primed individual | male | 43 | yes | 2025/5/28 | 62 | 68 | 99 | 91 | 58 |
| Wuhan-Hu-1 primed individual | female | 31 | yes | 2025/6/26 | 40 | 46 | 78 | 28 | 39 |
| Wuhan-Hu-1 primed individual | male | 23 | yes | 2025/6/26 | 67 | 65 | 92 | 25 | 188 |
| Wuhan-Hu-1 primed individual | female | 69 | yes | 2025/7/2 | 109 | 88 | 136 | 39 | 242 |
| Wuhan-Hu-1 primed individual | female | 11 | yes | 2025/7/2 | 81 | 64 | 74 | 18 | 58 |
| Wuhan-Hu-1 primed individual | female | 18 | yes | 2025/7/4 | 162 | 91 | 90 | 29 | 173 |
| Wuhan-Hu-1 primed individual | male | 43 | yes | 2025/7/11 | 13 | 19 | 29 | 123 | 67 |
| Wuhan-Hu-1 primed individual | male | 20 | yes | 2025/7/18 | 92 | 54 | 58 | 35 | 128 |
| Wuhan-Hu-1 primed individual | male | 57 | yes | 2025/7/24 | 69 | 26 | 43 | 80 | 276 |
| Wuhan-Hu-1 primed individual | female | 54 | yes | 2025/8/1 | 98 | 62 | 39 | 17 | 118 |
| Wuhan-Hu-1 primed individual | male | 26 | yes | 2025/8/1 | 92 | 40 | 55 | 25 | 220 |
| Wuhan-Hu-1 primed individual | female | 28 | yes | 2025/8/8 | 44 | 44 | 46 | 24 | 74 |
| Wuhan-Hu-1 primed individual | female | 25 | yes | 2025/8/8 | 68 | 78 | 112 | 47 | 48 |
| Wuhan-Hu-1 primed individual | female | 23 | yes | 2025/8/8 | 89 | 43 | 31 | 73 | 169 |
| Wuhan-Hu-1 primed individual | female | 39 | yes | 2025/8/8 | 112 | 55 | 47 | 66 | 305 |
| Wuhan-Hu-1 primed individual | female | 48 | yes | 2025/8/26 | 75 | 38 | 21 | 41 | 171 |
| Wuhan-Hu-1 primed individual | female | 37 | yes | 2025/8/26 | 59 | 64 | 61 | 19 | 66 |

**Figure S1.**
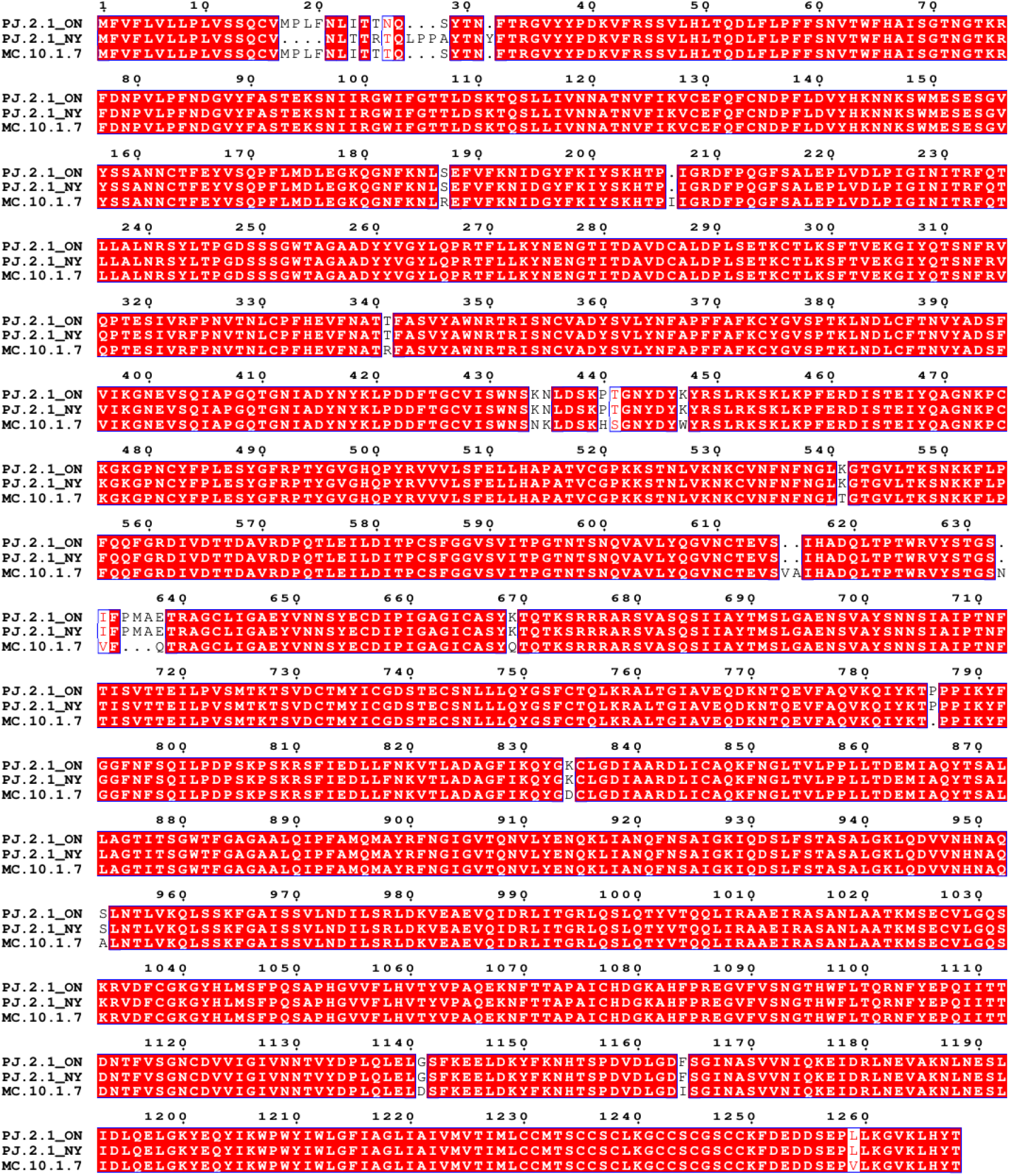
Amino acid sequence alignment of Spike glycoproteins from SARS-CoV-2 varian t PJ.2.1 isolates and MC.10.1.7. Multiple sequence alignment of full-length spike proteins from representative PJ.2.1 isolates identified in Ontario, Canada (PJ.2.1_ON) and New York, USA (PJ.2.1_NY) alongside the reference lineage MC.10.1.7. Sequences were aligned using MAFFT and visualized with ESPript 3.2^2^. Identical residues are highlighted with red backgrounds and white text; blue frames denote residues with similar physicochemical properties; dots represent alignment gaps.

**Figure S2.**
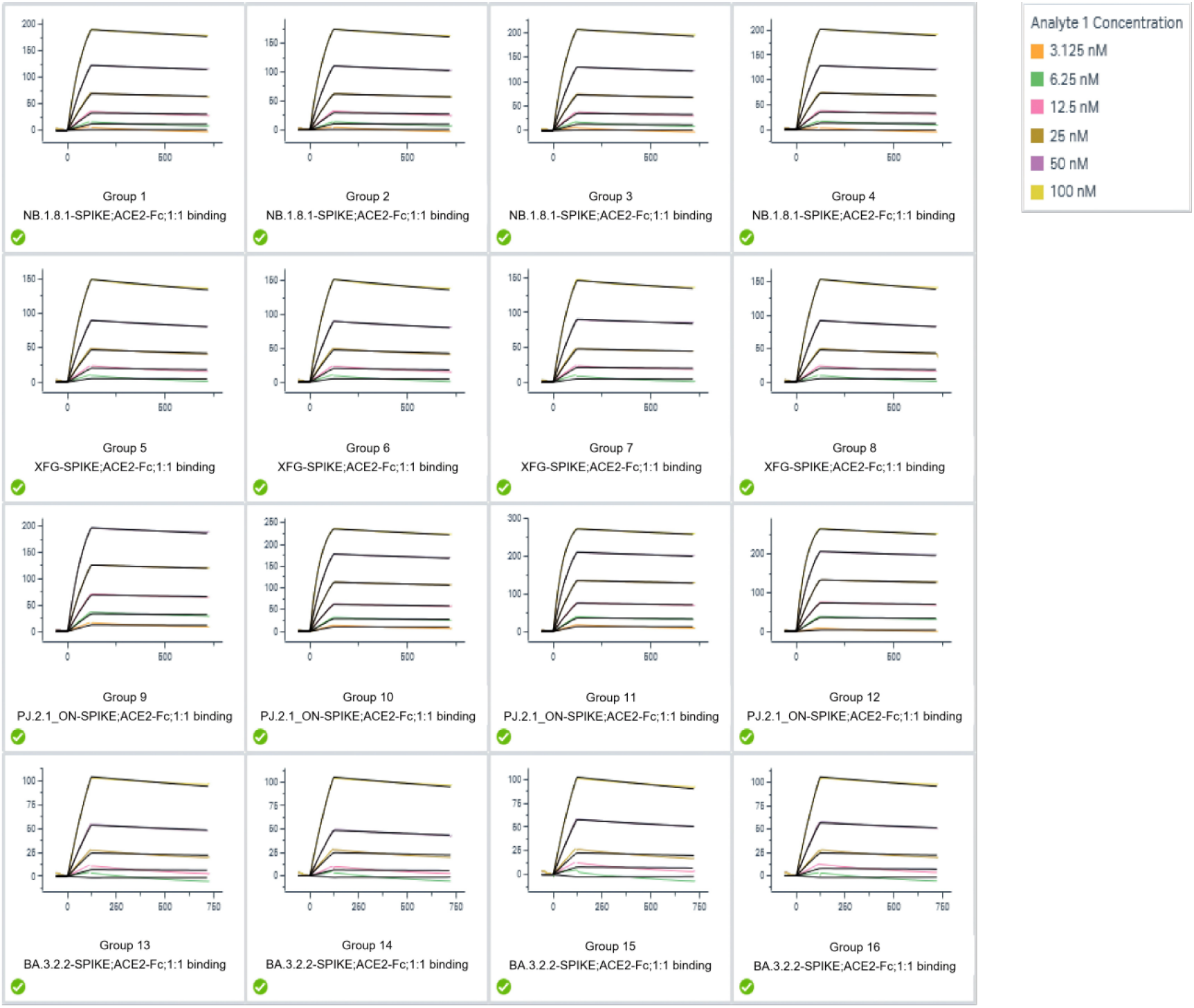
Surface plasmon resonance (SPR) sensorgrams measuring binding kinetics of human ACE2 to SARS-CoV-2 variant spike trimers. Kinetic profiles were acquired using multi-cycle kinetics runs on a Biacore 8K system. Twofold serial dilutions of purified SARS-CoV-2 spike glycoproteins were flowed over human ACE2 captured on Protein A sensor chips at room temperature. Raw binding responses are depicted as colored curves, overlaid with black lines denoting theoretical fits derived from a 1:1 binding model. Kinetic parameters, including association rate constant (ka), dissociation rate constant (kd), and equilibrium dissociation constant (KD), were analyzed using Biacore 8K Evaluation Software (version 4.0.8.20368).

